# Bioelectricity generation from live marine photosynthetic macroalgae

**DOI:** 10.1101/2021.04.16.440133

**Authors:** Yaniv Shlosberg, Nimrod Krupnik, Tünde N. Tóth, Ben Eichenbaum, Matan Meirovich, David Meiri, Omer Yehezkeli, Gadi Schuster, Álvaro Israel, Noam Adir

## Abstract

Conversion of solar energy into electrical current by photosynthetic organisms has the potential to produce clean energy. Previously reported bio-photoelectrochemical cells (BPECs) have utilized unicellular photosynthetic microorganisms. In this study, we describe for the first time BPECs that utilize intact live marine macroalgae (seaweeds) in natural seawater or saline buffer or natural seawater. The BPECs produce electrical currents from of >50mA/cm^2^, from both light-dependent (photosynthesis) and light independent processes. These values are significantly greater than the current densities that have been reported for single-cell microorganisms. The photocurrent is inhibited by the Photosystem II inhibitor DCMU, indicating that the source of light-driven electrons is from water oxidation via NADPH and other reduced molecules. We show here that intact seaweed cultures can be used in a large-scale BPEC containing seawater that produces bias-free photocurrent. The ability to produce bioelectricity from intact seaweeds may pave the way to future development of a low-cost energy technology using BPECs.

## INTRODUCTION

Photosynthesis is the process in which light energy is converted into storable chemical energy in the form of polycarbonic compounds, occurring in organisms that thrive in almost all environments that are accessible to light. Marine macroalgae, also known as seaweeds, have key ecological roles and are important primary producers in marine ecosystems. Seaweeds are taxonomically classified into 3 main groups: green (*Chlorophyta*), red (*Rhodophyta*) and brown (*Ochrophyta*)(*1*). While all seaweeds contain chloroplasts, they differ in their size, morphology, pigment types and light harvesting complexes (*2, 3*). Many seaweeds have a leaf-like sheets (*thallus*) or they may be filamentous or branched (*3*). Seaweeds are estimated to contribute up to 1 Pg C per year to global primary productivity (*4*). The photosynthetic efficiency of aquatic biomass is on average 6 to 8 % higher than that of the average photosynthetic efficiency of 1.8 to 2.2 % of terrestrial biomass (*5*). Seaweeds are mild halophiles as their seawater environment contains approximately 600mM salts (primarily NaCl).

The green seaweed *Ulva* sp. is abundant worldwide and proliferates seasonally along the Israeli Mediterranean Sea intertidal and rocky shores (*6*). In cultivation, *Ulva* species are reported to possess high growth rate of about 20% biomass gain or more per day (*7*–*9*). This high growth rate is correlated with high photosynthetic efficiency potential and electron transfer rate (ETR) (*10, 11*). The photosynthetic efficiency of *Ulva* may be attributed to its large surfaces to volume ratio, the coexistence of C3 and C4-like photosynthetic pathways (*9, 12*) and its efficient inorganic Carbon Concentrating Mechanism (CCM) based on HCO_3_^-^ uptake (*13*–*15*). The fast growth rate and tissue simplicity allows to easily obtain large amounts of plant biomass from marine seaweeds. In recent decades, attempts to utilize algal biomass as a source of bioenergy have been made. These include extraction of algal oils for biodiesel production, conversion of carbohydrates to hydrogen, bioethanol and biogas by means of hydrolyzation and fermentation(*16*–*18*). Yet attempts to use macroalgae as an efficient source of renewable and clean energy in a non-destructive manner have not yet been reported.

The production of electrical current by microbial fuel cells (MFCs) is to date far more mature (*19*–*25*). MFCs utilize the ability of bacteria to perform external electron transfer (EET) to the anode of an electrochemical cell (*26*) or to accept electrons from its cathode (*26, 27*). A derivative of MFCs are bio-photoelectrochemical cells (BPECs) that produced photocurrent from photosynthetic systems: isolated photosystems, thylakoid membranes or from whole microorganisms such as cyanobacteria and microalgae (*28*–*36*). One advantage of using photosynthetic microorganisms as a source of energy is their ability to remove atmospheric CO_2_ during photosynthesis, which makes them a less expensive and a potential clean source of energy that is beneficial to the environment. The power and stability of such BPECs depend on the photosynthetic material and the components of the electrochemical cell, including the need to add molecules that perform mediated electron transfer (MET). Untreated live cyanobacteria and microalgae have been shown to produce up to 10 µA/cm^2^/(mg chl) (*37*), while gentle treatment with a microfluidizer increased the current to 40 µA/cm^2^. Addition of quinones to live cells was shown to increase the photocurrent up to 60 µA/cm^2^. Higher photocurrents of up to 500 µA/cm^2^ were obtained by utilization of isolated thylakoid membranes from spinach with the addition ferricyanide ions (FeCN) for MET. In most cases, addition of the PSII inhibitor 3-(3,4-dichlorophenyl)-1,1-dimethylurea (DCMU) to photosynthetic organisms or isolated membranes inhibits harvested photocurrent, indicating that the major electron source is PSII (*30, 38, 39*) However, in some cases electrons from the respiratory system can be collected by the photosynthetic membrane, bypassing PSII (*37, 38, 40*). As isolated membranes are disconnected from photodamage cellular repair mechanisms, a decrease in current levels is apparent after 10 minutes of illumination (*30*). Recently, we have discovered that the main native electron mediator in cyanobacteria is NADPH (*40*), and that addition of exogenous NADP^+^ can significantly enhance the photocurrent of intact cells from 5 to 30 µA / cm^2^ / mg chl. In contrast to FeCN, NADPH is not toxic and, therefore, has the potential to be integrated in algae cultivation pools without harming the cells. Another method to increase the photocurrent production in BPEC is by utilization of cyanobacterial biofilms (*38, 41, 42*) which were reported to be able to perform direct electron transfer (DET) as opposed to the MET systems described above. Moreover, the tight arrangement in the biofilm increases the density of the cells to form a tissue-like structure and therefore increases the number of cells which may contact the interface of the electrode. Utilization of marine cyanobacteria in BPECs in high ionic strength electrolytic solutions, can improve the collected current by at least 2 orders of magnitude (*38*) due to increased conductivity (*43*).

In this work, we present for the first time a BPEC that can produce substantial electric current from seaweeds with promising future applications. We show that intact seaweeds can produce current in both dark and light as well as in a bias free macro system.

## RESULTS AND DISCUSSION

### Live *Ulva* produce an electric current in a bio-photo electrochemical cell

In our previous studies, we have described the harvesting of photocurrent from cyanobacteria (*37*) or isolated thylakoid membranes (*30, 44*) applied to the anode. The colloidal state of cells in solution physically form a layer on inorganic/metallic electrodes by sedimentation or by dipping the electrodes in the solution. This procedure is not suitable for utilization of seaweeds in BPECs, as they tend to float and have poor affinity to flat electrode materials. A first attempt to produce photocurrent from the seaweeds was done by designing a BPEC in which the *Ulva* is attached to different anode materials with a cover glass. The ability of carbon cloth, indium tin oxide, aluminium or stainless steel to function as the anode and produce photocurrent was evaluated. The highest photocurrent was obtained using stainless steel (Fig. S1).

Similar to previous work with thylakoids and cyanobacterial BPECs (*30, 44*), photocurrent requires close association between the *Ulva* tissue and the anode. However, the smooth texture of the *thallus* and their tendency to slide away (exacerbated by the oxygen bubbles that are released during photosynthesis), decreased the contact between the *Ulva* and the anode. We thus improved BPEC connectivity by using a standard stainless-steel clip that holds the *Ulva* tightly within the electrolyte solution. The metal clip is covered by plastic sheathing except for the surface in contact with the thallus. The interface area of the clip (both sides) between the clip and the *Ulva* was 0.08cm^2^. Platinum was used as the cathode and Ag/AgCl 3M NaCl as the reference electrode.

### Light drives increased electric current in intact *Ulva*

To study whether *Ulva* can produce current in dark or light, *Ulva* was placed in the BPEC and the current harvested by the anode was measured by chronoamperometry (CA) in 50 ml of a 0.5 M NaCl solution, with a bias potential of 0.5 V on the anode vs. Ag/AgCl 3M NaCl. This bias value chosen because it was the minimal value that produced maximal current. It was also the optimal value that was applied in our previous work with cyanobacterial driven BPECs (*40*). The illuminated area of the thallus was 0.5 cm^2^, as shown in in Fig. 1a and Fig. S2. To serve as a control material, we used a naturally bleached piece of *Ulva*, acquired from the same cultivation tank as the live green *Ulva*. Under continuous illumination with the intensity of 1 Sun, a maximal current density of ∼ 25 or ∼ 40 mA / cm^2^ was obtained after ∼10 min of measurement in dark or light, respectively (Fig. 1b). The current density that was harvested from the *Ulva* thallus in low salinity and without the addition of exogenous electron mediators is about three orders of magnitude larger than previously obtained by cyanobacterial based BPECs. This is a result of the combination of the high rate of photosynthesis and the enhanced conductivity available in the high ionic strength electrolyte solution (see below). As the clip-type anode covers the thallus, we postulate that light driven reactions in the cells that are adjacent to the anode induce a flux of reduced molecules that flow to the anode through the outer surface of the thallus. We thus wished to evaluate the *thallus* active area that contributes to the current production as function of its distance from the anode, in both dark and light. CA was performed in different sizes of *thallus* in both dark and light (Fig. S3). The measurements showed that when the *thallus* is cut so that it fully covered by the anode clips, a current density of about 15.6 mA/cm^2^ is obtained in either dark or light. When the *thallus* piece is larger, tissue becomes available for illumination resulting in higher currents, with light currents ∼2.6 times greater than in the dark. The current production was plotted against the maximal distance of the *thallus* from the anode (Fig. S3c). The maximal effective distance from the anode in both dark and light was determined to be ∼ 0.2 cm. Fitting analysis of the curves showed a logarithmic pattern (Fig. S3d) in both dark and light, indicating that the ability of the *thallus* to contribute for the current production decreases as function of distance from the anode. Thus, the active area of the thallus is ∼25% of the entire cutting. However, the current density values reported here are based on the size of the electrode surface (0.08cm^2^).

**Fig 1.**
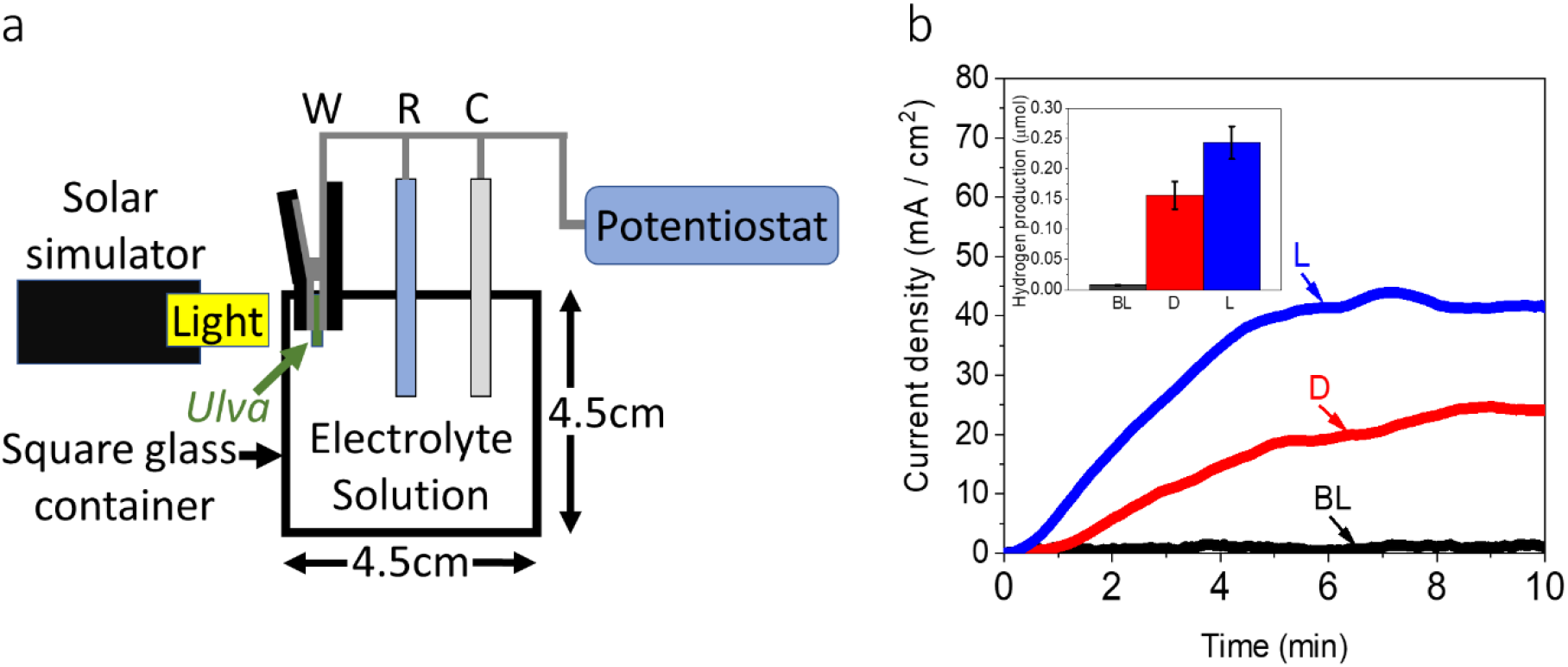
Description of the system and CA and Hydrogen production measurement. **a** schematic drawing of the measurement setup which is composed of a stainless-steel clip as anode, platinum counter electrode cathode and Ag/AgCl 3M NaCl reference electrode (RE) dipped in 50 mL electrolyte solution. A solar simulator is placed horizontally to illuminate a round *Ulva* leaf (diameter = 1 cm) with an intensity of 1 SUN. **b** CA measurements of bleached and green *Ulva* were measured in dark and light for 10 min. The onset of the light was at 0 min). CA of naturally bleached *Ulva* in light (BL, black), green *Ulva* in dark (D, red) and green *Ulva* in light (L, blue). Following the measurement, hydrogen production was quantified by GC. The inset shows hydrogen production after 10 min of bleached *Ulva* in light (BL, black), green *Ulva* in dark (D, red) and green *Ulva* in light (L, blue). The error bars represent the standard deviation over 3 independent measurements.

We have previously shown that current harvested from photosynthetic organisms at the anode results in significant proton reduction to molecular hydrogen on the Pt cathode, although other reduction reactions are possible due to the release of molecules from the *thallus*. To quantitate hydrogen production, the top of the glass box was sealed with a thick layer of parafilm. Immediately following the measurement, the gas phase above the solution was withdrawn by a syringe and its hydrogen content was quantitated by GC / MS (Fig. 1b). Hydrogen quantities of ∼ 0.15 or ∼ 0.25 µmol H_2_ were obtained after ∼10 min in dark or light, respectively.

### Higher salinity increases photocurrent from *Ulva*

The effect of the influence of high concentrations of ions on an electrolyte solution, especially a mixture such as found in natura seawater is highly complex (*45*). Increasing the ionic strength of a BPEC solution promotes increased conductivity (and reduction of the BPEC internal resistance) of the electrolyte (*46*). The benefits of the presence of high ionic strength electrolyte solutions have been previously shown for microbial fuel cells (*47, 48*). For most photosynthetic organisms (or their isolated components), a highly saline buffer is not experimentally feasible, as it induces stress on organisms or systems whose native habitat is fresh water. Since *Ulva* grows in seawater, and we are using the intact organism, the ionic strength of the surrounding electrolyte solution can be increased without harm to that of natural seawater. CA of *Ulva* under illumination was measured at increasing NaCl concentrations of 1 mM, 10 mM, 100 mM, 500 mM and natural seawater (∼600mM). The measured photocurrent significantly increased as a function of the NaCl concentration reaching maximal photocurrents of ∼2, 10, 30, 40 and 50 mA / cm^2^ respectively (Fig. 2). The possible influence of pH on the photocurrent production was also assessed. For this purpose, CA of *Ulva* was measured in MES buffer with 0.5 M NaCl with different pH values of 5, 6, 7 and 8 under illumination. A maximal photocurrent of ∼ 45 mA / cm^2^ was obtained for pH = 5. However, the differences in the photocurrent productions between all pH values was not significant as shown in Fig. S4.

**Fig. 2.**
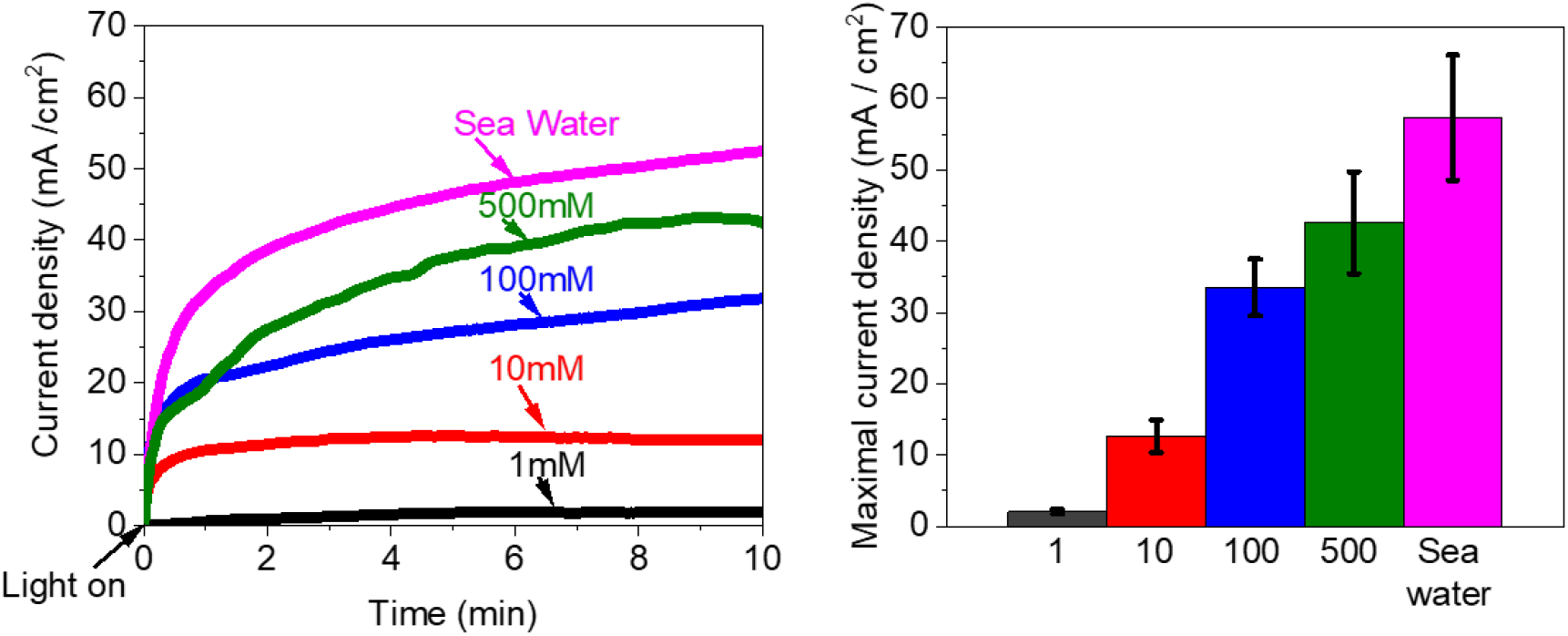
*Ulva* can produce a higher photocurrent at high salinity. CA of *Ulva* was measured in water containing increasing NaCl concentrations. **a** Representative CA measurements at increasing NaCl concentrations. 1 mM (black), 10 mM (red) and 100 mM (blue) and sea water (magenta) in light. **b** Quantitative analysis of maximal photocurrent production. 1 mM (black), 10 mM (red) and 100 mM (blue) and sea water (magenta). Error bars represent the standard deviation of three independent biological measurements.

### Endogenous and exogenous mediators can perform mediated electron transfer between *Ulva* and the anode

We have recently reported that under illumination, NADPH is the major endogenous mediator which transfer electrons between cyanobacterial cells and the anode (*40*). There, 2D-fluorescence maps (2D-FM) measured the accumulation of NAD(P)H in the external cellular media (ECM). As we hypothesized that MET is also the major mechanisms of electron transfer in macroalgae BPECs, CA of *Ulva* was performed in 0.5 M NaCl for 10 min followed by 2D-FM measurement of the ECM (Fig. 3). The strong peak at λ_max_(ex) = 350/ λ_max_(em) = 450 nm that is the major fingerprint of NAD(P)H (*40, 49*) was clearly identified. The concentration of NAD(P)H in the ECM was quantified(*40*) and determined to be ∼0.015 µM. An additional possible electron donor is hydroxyl ions that are naturally secreted by seaweeds to regulate the pH at the *thallus* surface (*50*).

**Fig. 3.**
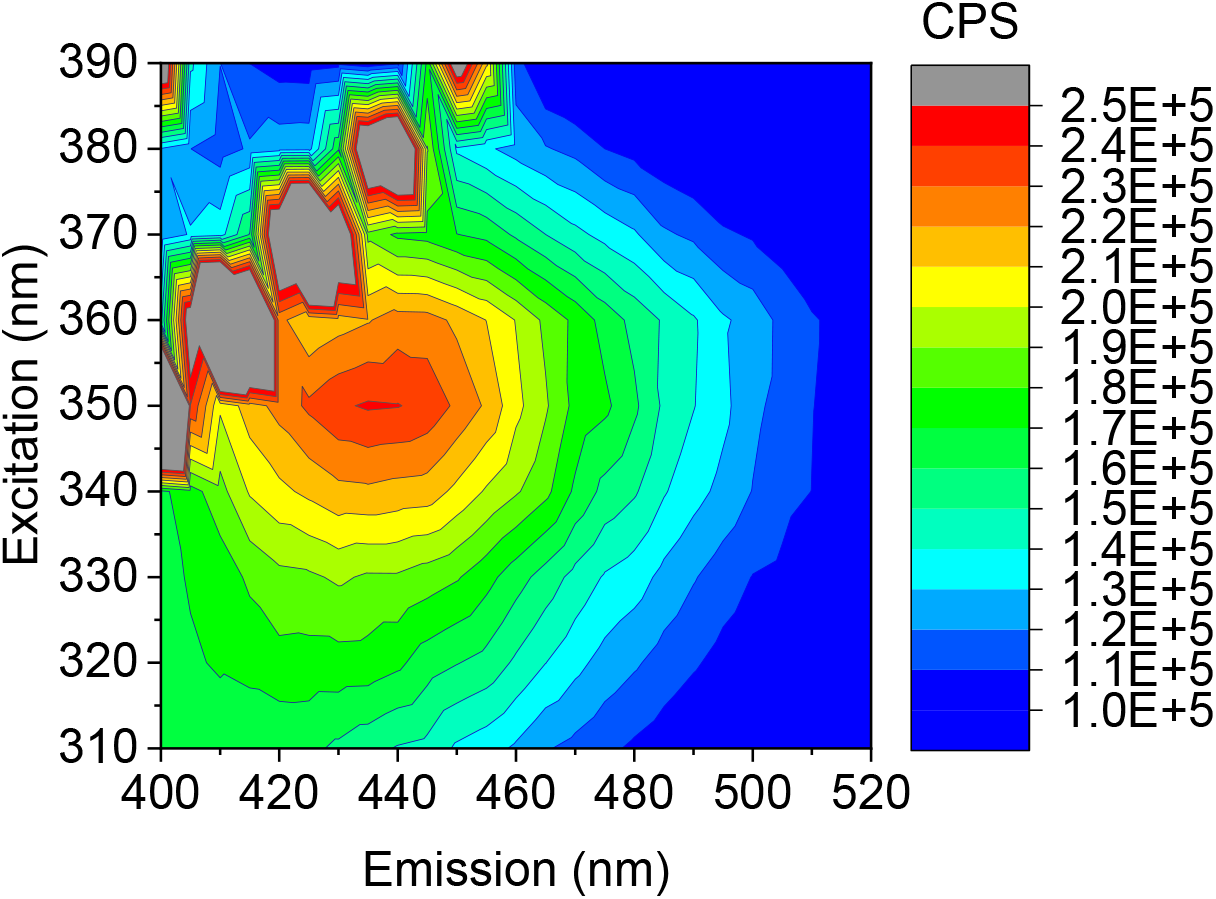
*Ulva* secretes NAD(P)H to the external cellular media. 2D-FM spectra of the ECM of *Ulva*. The obtained peak at (λ_max_ (ex) = 350, λ_max_ (em) = 450 nm) correspond to the spectral fingerprint of NAD(P)H (*40*). The lines of diagonal spots that appear in all of the maps presented here and in the following figures results from light scattering of the Xenon lamp and Raman scattering of the water (*60*).

An additional way to improve current harvesting is the addition of exogenous mediating molecules. Previous studies reporting bioelectricity production from photosynthetic and non – photosynthetic microrganisms have utilized FeCN as a soluble electron mediator for MET between the microorganisms and anodes (*51*–*55*). CA of *Ulva* in water containing 0.5 M NaCl was measured in the presence of 1 mM FeCN. Addition of FeCN increased the measured photocurrent 2-fold after 10 min (Fig. 4a). We determined that the role of FeCN is as an exterior mediator as *Ulva*, incubated for 30 min in the presence or absence of 5 mM FeCN in DDW, did not show the presence of internalized FeCN (Fig. 4b).

**Fig. 4.**
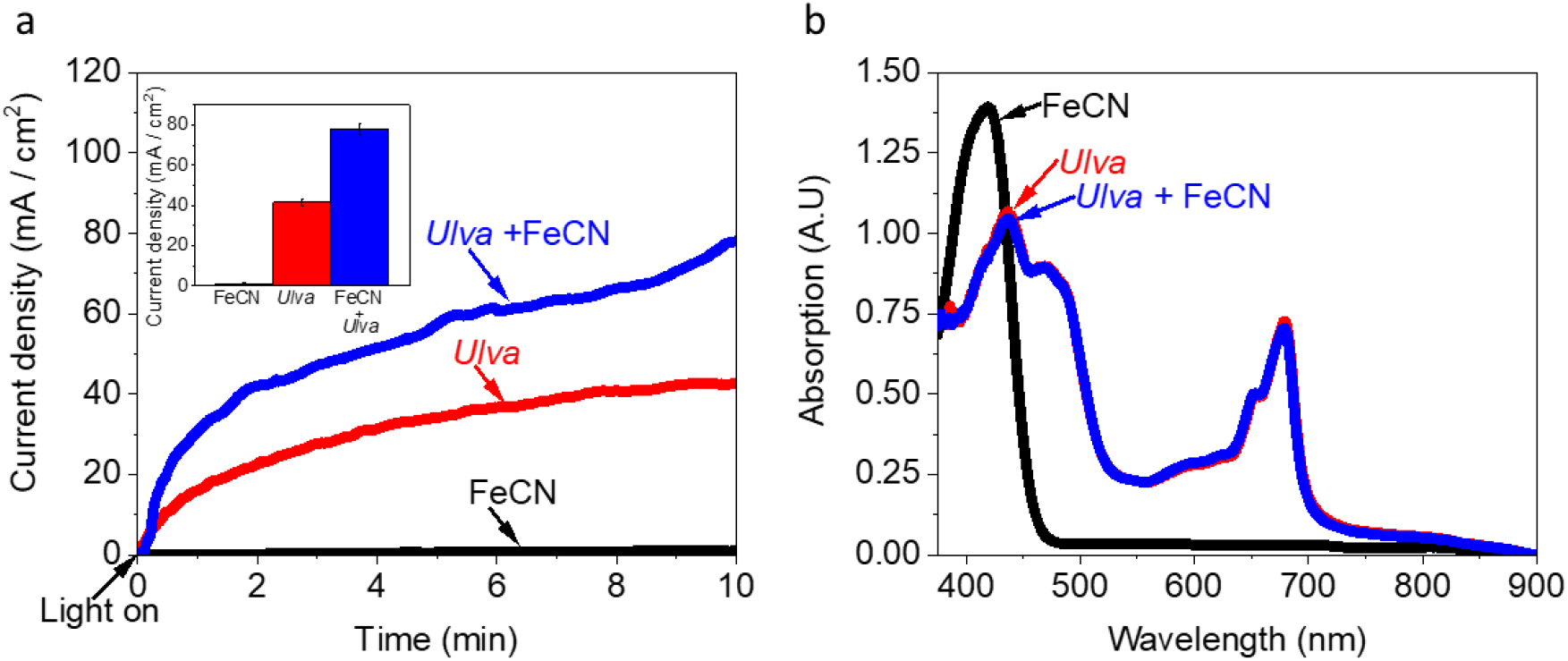
Ferricyanide ions mediate electrons between *Ulva* and the anode and increases the photocurrent. **a** CA measurement in 0.5 M NaCl solution of *Ulva* (red), *Ulva* + 1mM FeCN (blue) and 1mM FeCN without *Ulva* (black). The error bars in the inset represent the standard deviation of the maximal photocurrent over 3 independent measurements. **b** Absorption spectra of pure FeCN solution (2.5 mM) (black), *Ulva* after 0.5 h in DDW (red) and *Ulva* after 0.5 h in 5 mM FeCN (blue).

### The source of electrons harvested from illuminated *Ulva* is from water oxidation by PSII

Previous studies about current production from spinach thylakoid membranes reported that addition of the herbicide DCMU, abrogated photocurrent production (*30*). In the case of *Ulva*, addition of 100 μM DCMU to the BPEC electrolyte solution indeed inhibited photocurrent harvesting (Fig. 5a) indicating that PSII is the major source of photocurrent. To validate the inhibition of PSII by DCMU, a dissolved oxygen (DO) sensor was used to measure oxygen production of *Ulva* in the same experimental setup of the CA measurements in the presence or absence of DCMU (Fig. 5b and Fig. 6). The oxygen concentrations after 10 min of illumination in the presence or absence of DCMU were determined to be ∼ 0.05 and 4 mg / mL respectively. These results imply that under illumination most of the photocurrent derives from the photosynthetic pathway initiated by PSII, however the involvement of NADPH as shown above indicates that both photosystems are involved in photocurrent production.

**Fig. 5.**
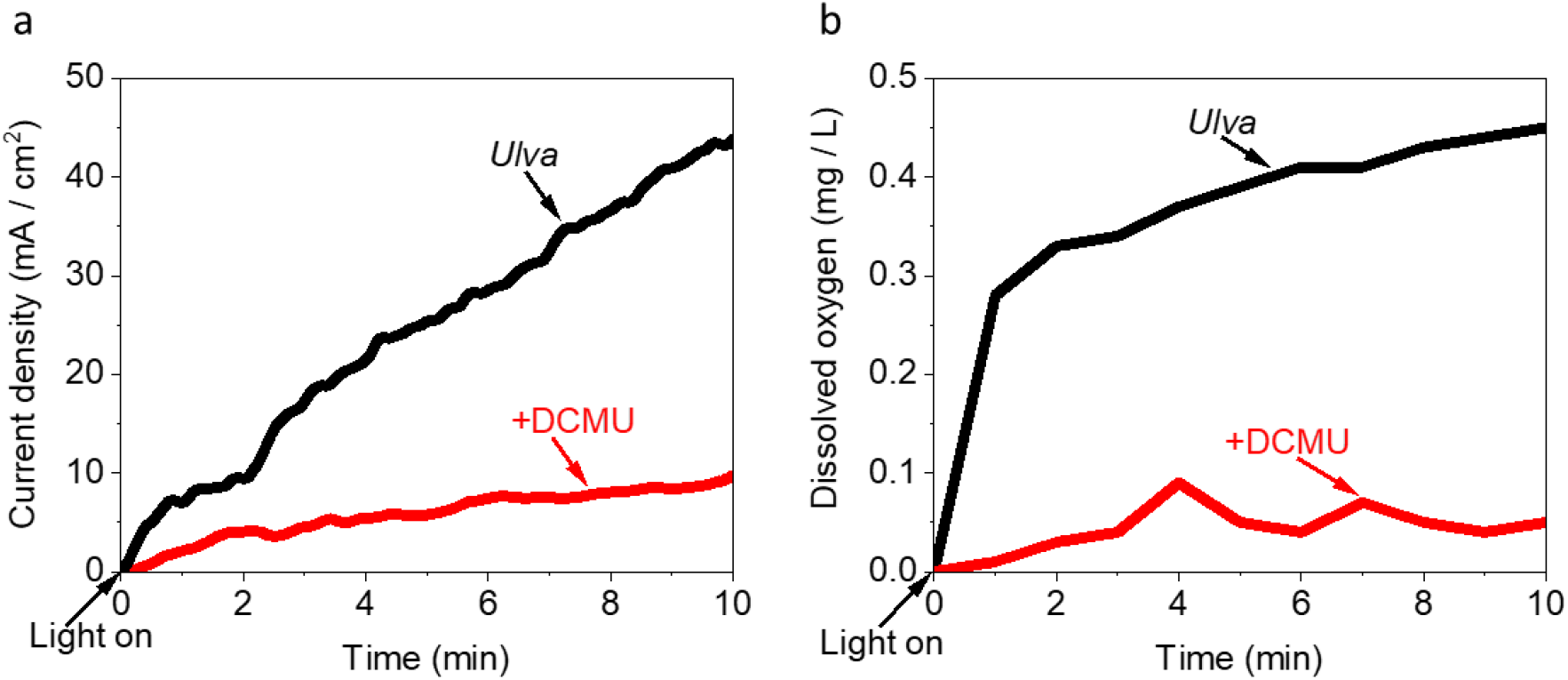
DCMU inhibits the photocurrent production of *Ulva*. CA and DO of *Ulva* were measured in under illumination with and without addition of 100 µM DCMU. **a** CA of *Ulva* (black), CA of *Ulva* + 100 µM DCMU (red). **b** DO measurements of *Ulva* (black), and *Ulva* + 100 µM DCMU (red).

**Fig. 6.**
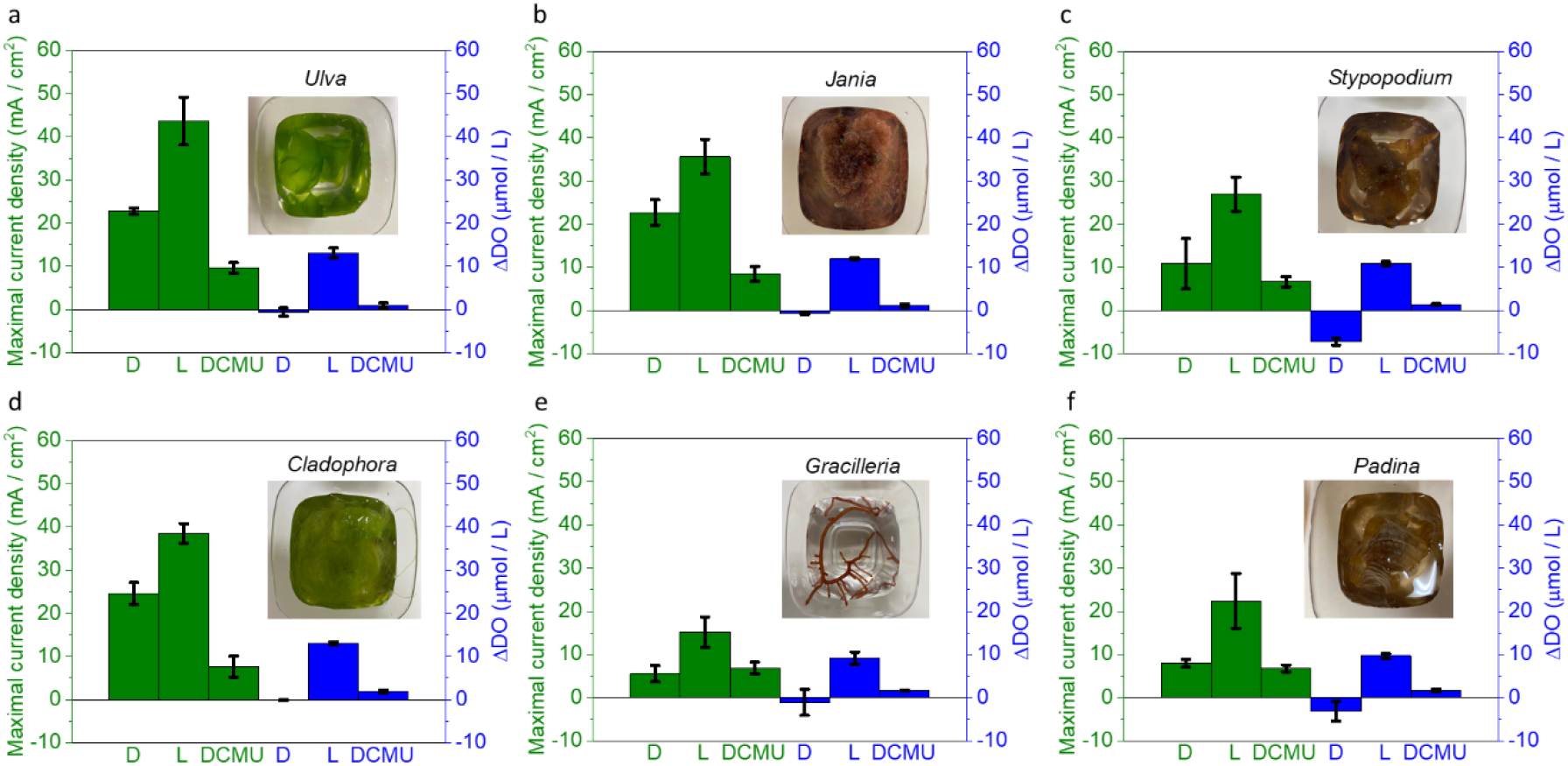
Photocurrent and DO measurements of seaweeds from various taxonomic groups. CA and DO were measured for the 6 different seaweeds in dark, light and in light + 100 µM DCMU. CA and DO measurements of **a** *Ulva*, **b** *Jania*, **c** *Stypopodium*, **d** *Cladophora*, **e** *Gracilaria* and **f** *Padina*. In all panels the left 3 green represents the current density after 10 min and the right blue Y axis represents the change in DO quantities after 10 min. The error bars represent the standard deviation over 3 independent measurements. The names and photos of the measured seaweeds are displayed in the panels.

The kinetics of the two reactions, oxygen evolution and electron harvesting by the BPEC are not similar. Oxygen evolution initiates (Fig. 5b) and ceases relatively quickly (seconds) upon initiation and termination of illumination. During this period, the concentration of reduced molecules (especially NADPH) within the *thallus* increases and diffuses towards the anode clips. We have previously shown that operation of the BPEC increases the movement of NADPH towards the anode, increasing the measured current (*40*), and this is most likely the case here as well. Following termination of the illumination, the BPEC continues to collect electrons until the current returns to the dark value. We have previously shown that in cyanobacteria, addition of external glucose supports oxidative phosphorylation which extends the ability of the system to continue to provide electrons to the BPEC (*37, 40*). In the system presented here, the *Ulva* tissue continues to efficiently incorporate CO_2_ into glucose by the Calvin cycle (*9*). This internal glucose source can be transported to the mitochondria, where it produces a constant flow of NADH/NADPH which is exported to the cells and to the extracellular spaces. Thus, the current obtained during illumination is a combination of direct and indirect production of reduced molecules, leading to the very high currents reported here.

### Photocurrent and DO measurements of seaweeds from various taxonomic groups

The ability of *Ulva* to produce photocurrent in the BPEC raised the question whether this ability is unique to *Ulva* or whether it exists in other seaweeds. To address this question, we collected environmental samples of red algal seaweeds *Gracilaria* and *Jania*, brown algal seaweeds *Padina* and *Stypopodium* and green seaweed *Cladophora* from the Eastern Mediterranean coast of Israel (Fig. 6 and S5). We performed CA measurements on all species as described for *Ulva*. All seaweeds successfully produced current, with maximal values of ∼ 5 – 20, ∼ 15 – 45, ∼ 5 – 10 mA / cm^2^ in dark, light or light + DCMU, respectively. DO measurements showed a similar pattern for all seaweeds in which the difference DO concentration was ∼ -0.4 – 0, 0.4 – 0.6 and 0 – 0.05 mg / L in dark, light and in light + DCMU, respectively (Fig. 6). Under illumination the green seaweeds *Ulva, Cladophora* and the red alga *Jania* produced photocurrents 2-2.7 times higher than the current produced from the brown seaweeds *Stypopodium* and *Padina* or *Gracilaria*. Despite the differences in the measured values, it is possible that other factors rather than photosynthetic properties influence the photocurrent production such as freshness and texture of the seaweeds. *Ulva* and *Gracilaria* can be successfully cultivated in tanks provided with continuous seawater and aeration. On the other hand, *Jania, Stypopodium, Padina and Cladophora* are difficult to cultivate. Therefore, they were collected from native coastal rocks. The different environments affect the physiology of the seaweeds and as a result may also affect their ability to produce photocurrent. Moreover, *Stypopodium, Padina*, and *Ulva* thalli have a smooth texture while *Jania* and *Cladophora* are composed of small fibres and are more adhesive. *Gracilaria* has a bulker smooth texture whose attachment to the anode was poorer than the other species.

The samples described here were obtained either from growth vats using natural seawater circulation (*Ulva*) or from environmental samples. Environmental samples most likely have a small amount of bacteria growing on their surface, including photosynthetic bacteria (*56*).However the concentration of bacteria are much lower that that used in previously described cyanobacterial BPECs (*37, 40*) and thus are not the major source of current harvested from the macroalgae.

### Toward applicative technologies for electric current production from seaweeds

The ability to produce current in the native habitat of *Ulva* is a significant advantage for clean energy production as it allows the organisms to simultaneously grow and produce current and benefit from the high electrolyte conductivity of the seawater. Typically, BPEC technologies utilize a potential bias on the anode to improve the current production. Such process entails an extra investment of energy which in some cases is higher than the energy produced by the BPEC itself. *Ulva* is classically cultivated for the production of biomass used in the food and cosmetics industries, and more recently explored for the production of biofuel (*8, 57*). From a practical perspective, we aimed this work to demonstrate the potential of integrating an electric current production system directly during seaweed cultivation. CA was measured on-site during 5 h in the *Ulva* cultivation tanks. The lower half part of a round aluminium plate was attached to the tanks and used as anode. A platinum wire was used as cathode (Fig. S6). Intact *Ulva* were continuously moving in the water stream, associating with the anode, producing the electrical current. In order to simulate a scenario in which the BPEC is producing electrical current without being aided by external energy, the measurement was done bias free with a 2-electrode mode. Natural sunlight irradiance was used (and thus is fluctuating) and measured at the seawater surface to be as high as 200 µE / m^2^ s^-1^. The seaweeds in the cultivation pool produced a maximal current of ∼0.5 mA after 5 hr (Fig. 7). No significant current was produced in the absence of seaweeds while in the absence of photosynthesis (dark), about 20% of the current density continued to be collected. The results demonstrate the possibility of integrating a BPEC system directly in seaweeds cultivation pools showing that a significant electrical current may be continuously generated in such simple systems. When CA of *Ulva* was measured in the laboratory set up (in seawater) using the potentiostat in 2-electrode mode, a photocurrent of 1 µA / cm^2^ was obtained (Fig. S7).

**Fig. 7.**
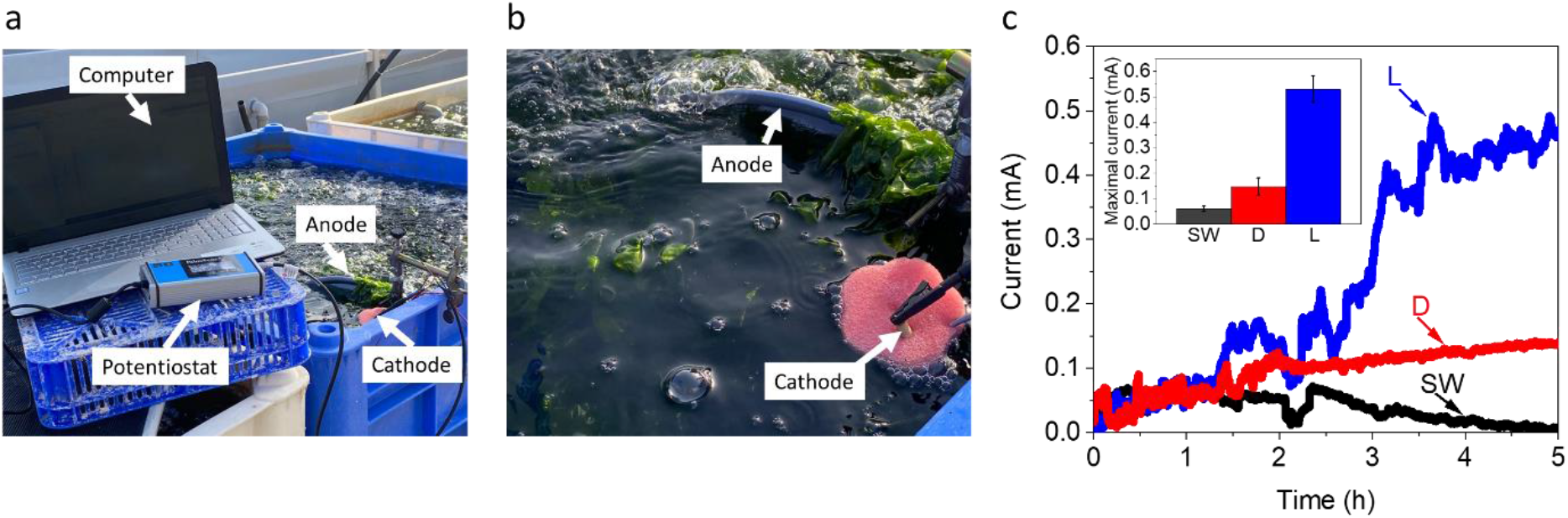
Toward applicative technologies for current production from seaweeds. Bias free current production of *Ulva* in its cultivation pool. The system is composed of a round Aluminium plate anode and a Platinum wire cathode which are dipped inside a cultivation pool with seawater and *Ulva*. The anode is held by a clamp and the cathode is placed in a sponge which is floating on the water surface. The pool is located on the seashore and contains a pipeline system which continuously stream water inside and outside of the pool. An average sunlight intensity of ∼200 µE / m^2^ was measured on at the pool surface. Pieces of *Ulva* are drifting in the water stream, hit the anode and produce electrical current. **a** A picture of the system. White arrows label the components of the system including the computer that operates the potentiostat, the potentiostat, the anode and the cathode. **b** A picture of the system which is focused on the inner part of the cultivation pool, the anode and the cathode. White arrows label anode and the cathode. **c**, Bias free CA measurements in the cultivation pool in seawater without *Ulva* under the sunlight (SW, black), with *Ulva* in dark (D, red) and with *Ulva* illuminated by natural (fluctuating) sunlight (L, blue) over 5 h. The inset displays the average maximal obtained current over 5 h. The error bars represent the standard deviation over 3 independent measurements.

### Possible external electron transport mechanisms in seaweeds based BPECs

The mechanisms of the electrochemical interactions between the seaweeds and the electrochemical cell are very challenging to understand. In fact, very little is known about their natural external electron transport mechanisms. Moreover, their association with the electrochemical components may modify the chemical characteristics of the *Ulva* surface, as we previously reported for cyanobacteria (*40*). In light of the results presented here, we suggest a model with several possible options that are based on the anatomy of the seaweed *Ulva*, known metabolic reactions and previous models which were reported for BPECs that were based on non – photosynthetic bacteria, cyanobacteria and microalgae (Fig. 8). The *thallus* of *Ulva* may have different sizes and shapes which are formed by different arrangements and densities of its cells. However, the sheet-like structure of the *thallus* exposes all cells to the interface with the seawater and in this way allowing them to perform different molecular export or import reactions with their environment (*6*). One possible mechanism for light dependent EET originates in the photosynthetic pathway which initiates in PSII which converts the sunlight into electric current and ends in reduction of NADP^+^ to NADPH by PSI. A fraction of the NADPH molecules may be transported to edge of the cell wall and from there to reduce the anode or to reduce an exogenous mediator, such as FeCN as suggested for eukaryote cells by Rawson et al. (*58*). The release of NADPH from the *thallus* may be enhanced when *Ulva* is associated with the anode of the electrochemical cell as previously reported for cyanobacteria (*40*). As shown above, transport of NADPH (or other potential mediators) can occur over a short distance (∼0.2 cm) within the *thallus*. As seaweeds also produce current which is not light dependant, NADPH may be produced in mitochondria, and then exported from the cells for intracellular transport. As *Ulva* and many other seaweeds secrete hydroxide ions in order to regulate the pH levels at their surface (*50*), we suggest that a high local concentration of hydroxide ions may also reduce the anode as occurs in alkaline water electrolysis (*59*).

**Fig. 8.**
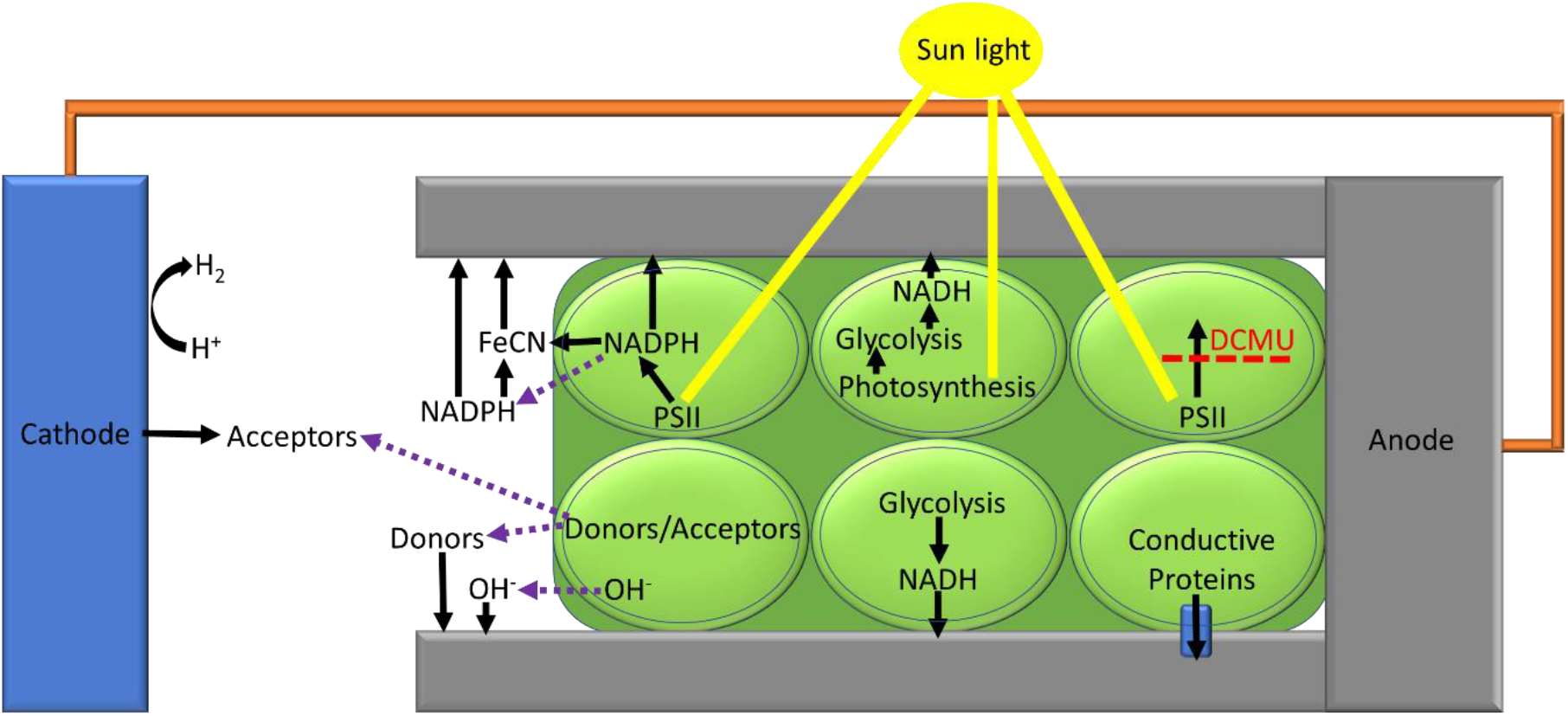
Possible external electron transport mechanisms in seaweeds based BPECs. Based on our findings and together with previous models which were reported for BPECs based on microalgae, cyanobacteria, non-photosynthetic bacteria, and thylakoid membranes, we propose a model for various possible EET mechanisms for the seaweeds based BPEC. The *Ulva thallus* is marked in dark green, and its cells are marked with round light green shapes. The sunlight is marked in yellow. The anode clip is marked in grey and the Pt cathode in a blue rectangular shape. A connective spring between the anode and cathode is marked in orange. The upper 3 cells of the *Ulva* describe EET mechanisms which are light dependent. The lower 3 cells of the *Ulva* describe EET mechanisms which are light independent. A small blue cone cylinder which is located in the lower right cell indicates a hypothetical membrane bound conductive complex. Labels indicate the different materials. Black arrows indicate the direction of potential electron transport. Purple dashed arrows indicate molecular secretion from the inner part of the *Ulva* cells to the ECM. A dashed red line which crosses a black arrow indicates the inhibition of the electron transport by DCMU.

## Conclusions

The potential for future clean bio(photo)energy technologies can be more easily met by avoiding the use of precious arable land, fresh water and fertilizers. For this reason, the use of seaweeds is optimal. This study shows for the first time that photocurrent can be harvested in a BPEC based on live seaweeds. We show that (1) seaweeds can produce biocurrent using simple and inexpensive metallic anodes that is more than 3 orders of magnitude higher than that obtained for microorganisms in fresh water-based buffers, (2) the biocurrent is available both in the dark and it is enhanced further in the light, with or without added bias on the anode, (3) the biological material (biomass) can be streamed down for use by the food, chemical or energy (biofuels) industries. This work paves the way for future developments of novel bio-electrochemical cells based on bulk photosynthetic organisms.

## Materials and Methods

### Seaweed cultivation, sampling and sample preparation

*Ulva* and *Gracilaria* thalli stocks were cultivated in land-based culture tanks as described in Israel et al. (*8*). *Jania, Stypopodium, Padina* and *Cladophora* were collected from the intertidal zone in Achziv and Habonim field sites on the eastern Mediterranean Sea coast. The sheet-like seaweeds *Ulva, Padina* and *Stypopodium* were cut down to 1 cm in diameter discs. *Jania, Cladophora* and *Gracilaria*, which all have more complex structures, were cut to an equivalent area of ∼ 0.79 cm^2^.

### Chronoamperometry measurements

#### a. Indoor CA measurements

All indoor measurements were done in 4.5 cm^3^ rectangular transparent glass vessels. The light source was produced using a solar simulator (Abet, AM1.5G) placed horizontally to illuminate the seaweeds with a solar intensity of 1 Sun (1000 W/m^2^). Determination of the solar intensity at the surface of the seaweeds was done as function of distance from the light source in an empty vessel neglecting small intensity losses caused by the glass and ∼ 0.5 cm of the electrolyte solution. bias free measurements were measured in 2 electrodes mode without application of electrical potential on the anode using the stainless-steel clip as WE and a Pt wire as CE in native sea water. All other indoor measurements were done in 3-electrode mode using the stainless-steel clip as WE, a Pt wire as CE and Ag/AgCl 3M NaCl as RE (in 3 M NaCl solution) (RE-1B, CH Instruments, USA) with an applied electric potential of 0.5 V on the anode in 0.5 M NaCl (except for the increasing salinity experiment described in Fig. 2 in the main text). In all measurements the current density was calculated based on the contact area between the WE and the seaweeds of 0.08 cm^2^.

#### b. Direct CA measurements from seaweeds cultivation tanks

CA measurements were done directly from the pools using the lower part of an aluminium plate as the WE, a Pt wire as the CE and Ag/AgCl 3M NaCl as the RE without added bias on the WE, under natural sunlight. Light intensity was measured by a light meter at the pool surface height.

### Dissolved oxygen measurements

DO measurements were performed using a DO meter probe (Hanna Instruments, HI-5421 research grade DO and BOD bench meter). The measurements were performed in the same experimental setup as the CA measurements. The DO probe was inserted into the electrolyte solution and the top of the glass container was sealed tightly with multiple layers of parafilm. A small magnetic bar was used to stir the electrolyte solution.

### Absorption measurements

Absorption spectra of *Ulva* were measured using a Shimadzu (UV-1800) spectrophotometer in 1 cm pathlength square cuvettes. For *Ulva* measurements the cuvette was filled with DDW. A rectangular piece of *Ulva* was cut and tightly attached to the inner sidewall of the cuvette which closer to the detector.

### H_2_ determination

The top of the BPEC was sealed with a thick layer of parafilm. CA measurements were conducted for 10 min. 1 mL of gas sample was taken from the glass vessel headspace using a syringe. Then the sample was injected into 1.8 mL glass sealed vials with screw caps suitable for auto-sampler injection (La-Pha-Pack). H_2_ production was determined by injecting a 50 µL sample into a gas chromatograph system with thermal conductivity detector (GC-TCD, Agilent 8860) equipped with a 5-Å column (Agilent, 25m x 0.25mm x 30μm).

## Acknowledgements

Funding was provided by a “Nevet” grant from the Grand Technion Energy Program (GTEP) and a Technion VPR Berman Grant for Energy Research. Some of the results reported in this work were obtained using central facilities at the Technion’s Hydrogen Technologies Research Laboratory (HTRL) supported by the Nancy & Stephen Grand Technion Energy Program (GTEP), the ADELIS Foundation and the Solar Fuels I-CORE. We thank Dr. Yifat Nakibly and Dr. Rachel Edrei for technical support. We thank Benjamin Eichenbaum, Itamar Shaul Eidelberg, Shaked Tzaban, Lee Keysar, Sonya Copperstein and for their technical assistance. Yaniv Shlosberg and Tünde N. Tóth are supported by fellowships of the Nancy & Stephen Grand Technion Energy Program (GTEP) and by a Schulich Graduate fellowship.

## Author contributions

YS and NA conceived the idea. YS, NK, AI, GS and NA designed the experiments. YS performed the main experiments. TNT, BE, OY and MM assisted in performing parts of different experiments. YS, NA, AI and NA wrote the paper. NA and GS provided all funding for the project. NA supervised the entire research project.

## Supplementary Materials

**Fig. S1.**
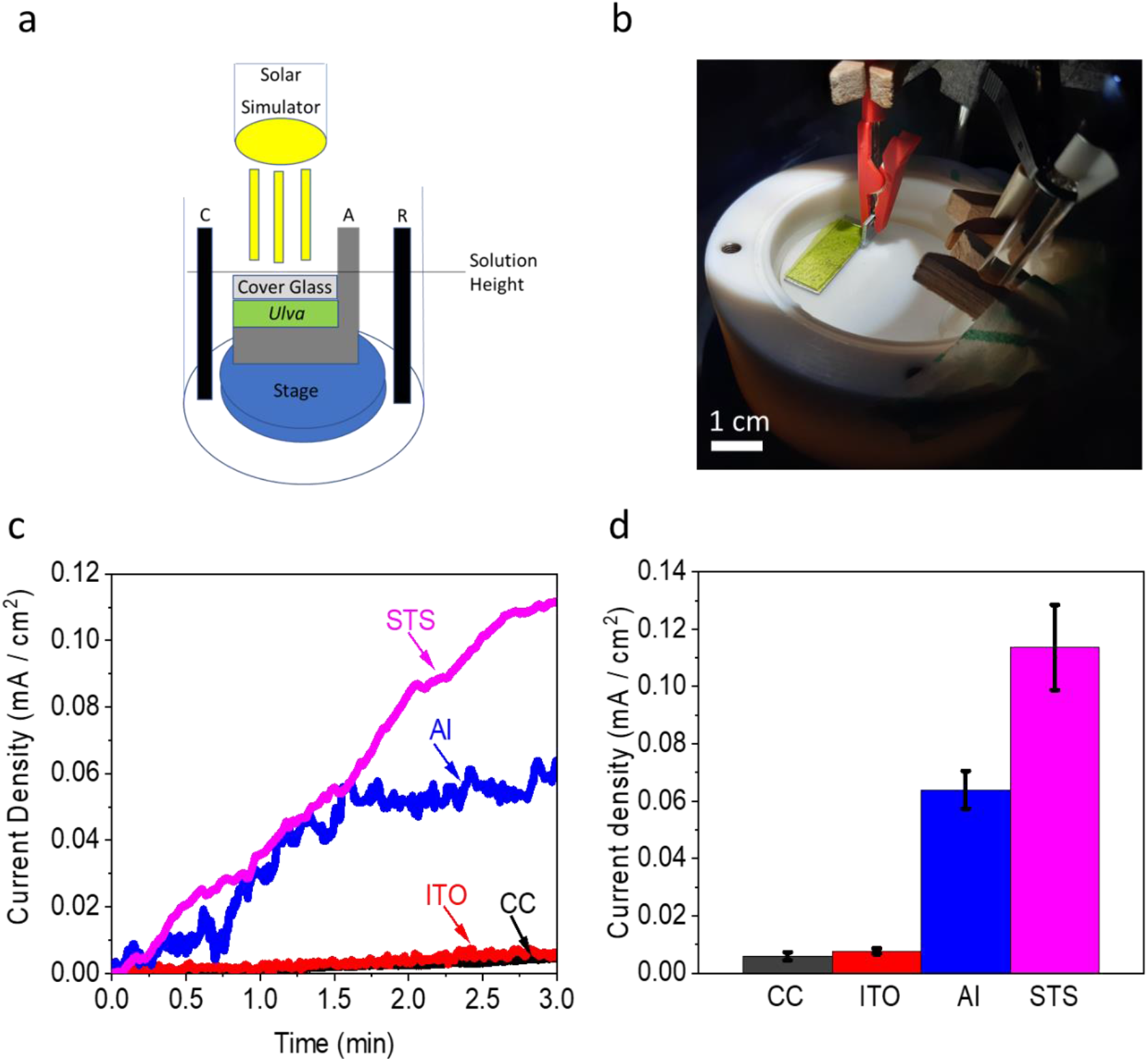
Utilization of different anodes materials for photocurrent production from *Ulva* in a bio-photo electrochemical cell. In order to integrate *Ulva* in a bio-photo electrochemical system a new BPEC design was built in which *Ulva* is placed above the anode and below a cover glass. The anode is placed on a stage and has a small part (0.5 cm^2^) which is bent upward with a 90^0^ angle. This part is connected to the potentiostat through a clip above the solution surface. Platinum cathode and Ag/AgCl NaCl 3M reference electrode are placed inside in 0.5 M NaCl solution. Illumination was done from above at intensity of 1 SUN (measured at the solution surface). **a**, Schematic drawing of the system. The anode, cathode and reference electrodes are marked as A, C and R respectively. **b**, Photo of the system under illumination. **c**, Chronoamperometry measurements of *Ulva* utilizing different material as Anode: Carbon cloth (CC, black), Indium tin oxide (ITO, red), Aluminium (Al, blue) and stainless steel (STS, magenta). **d**, maximal current production produced by the different anodes. Aluminium (Al, blue) and stainless steel (STS, magenta). The error bars represent the standard deviation over 3 independent measurements.

**Fig. S2.**
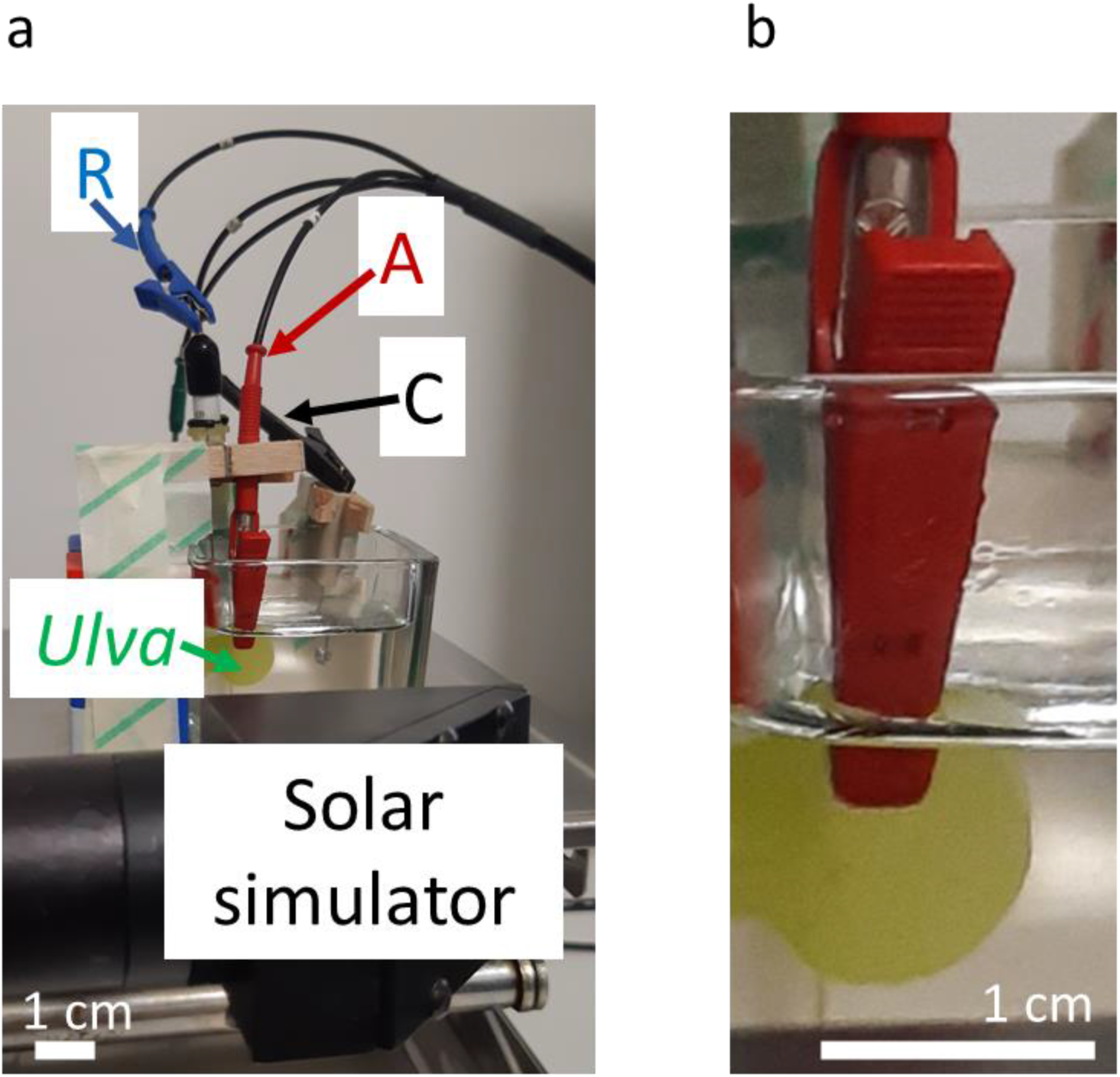
A photo of the system. labels and arrows point at the electric connections to the anode (red), cathode (black) and RE (blue). A green label and arrow point at the *Ulva*. A solar simulator label (black) shows the head of the solar simulator which illuminate the sample. **b**, an enlargement of panel **a** which focuses on the connection between the anode and the *Ulva*.

**Fig. S3.**
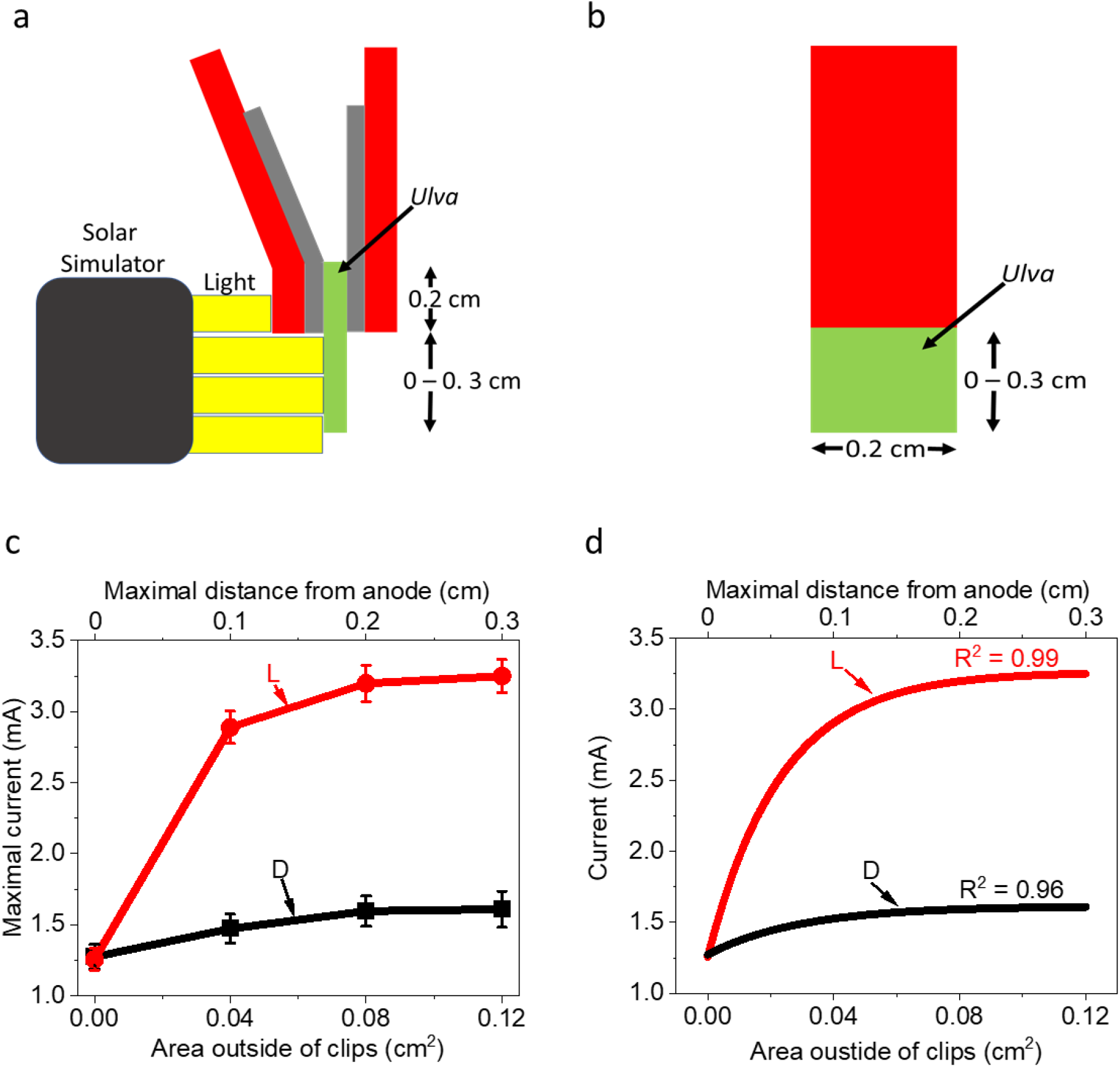
Evaluation of the maximal effective distance of the *thallus* from the anode clips. CA was measured in dark and light for 10 min. Different pieces of *thallus* with a fix width of 0.2 cm and variable increasing lengths of 0.2 – 0.5 cm. In all of the measurements the upper part of the *thallus* (0.2 cm) was fully covered by the anode clips. **a** schematic illustration of the system from a side view. Upon light illumination, only areas of the *thallus* outside of the clips were exposed to light. **b** schematic illustration of the system from a front view. The fixed and variable dimensions of the electrode (clip) area) and additional *thallus* area are denoted by arrows. **c** CA measurements of *thallus* with different sizes in dark (black) and light (red). The Y-axis represents the maximal current after 10 min (without normalization to the anode area). The lower X-axis represents the area of the *thallus* that is not covered by the anode clips. The upper X-axis represents the maximal distance of the *thallus* from the anode. The obtained current densities are displayed without any normalization to the electrode or *Thallus* area. The error bars represent the standard deviation over 6 independent measurements. **d** Exponential fitting of the CA measurements (shown in panel C) in dark (black) and light (red). R^2^ values are indicated above each curve.

**Fig. S4.**
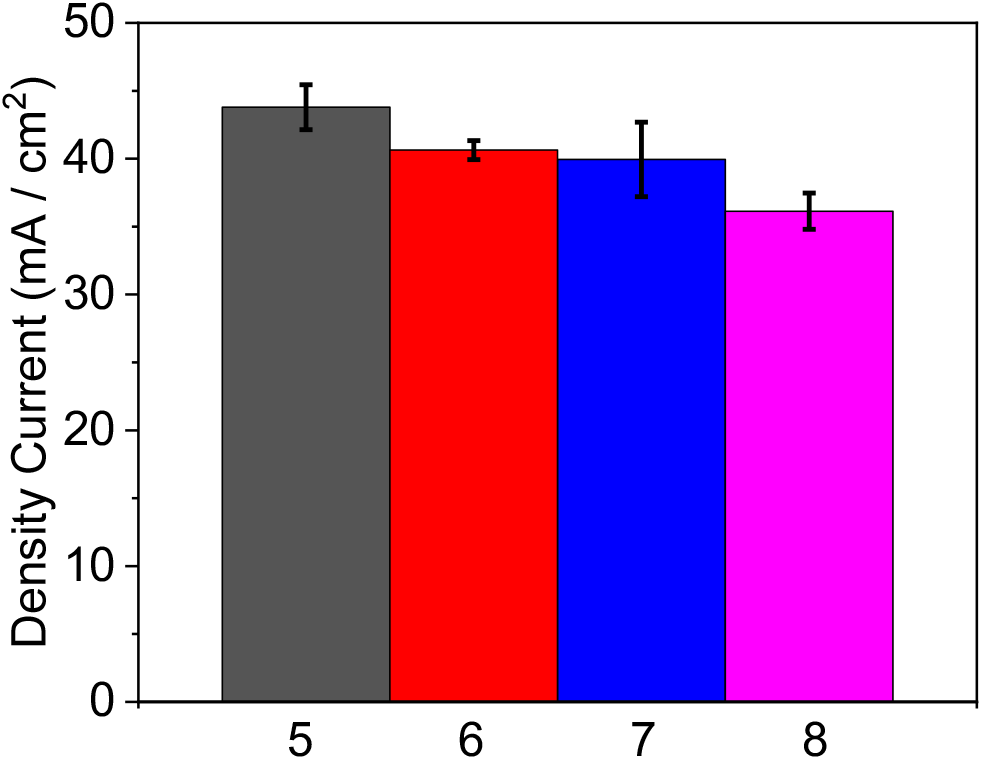
The pH of the electrolyte has a small effect on the current production. CA of *Ulva* was measured in light in solutions of MES buffer with 0.5M NaCl with increasing pH values (5 – 8). The maximal current production was obtained for pH = 5, however, it was not significantly higher than the solutions with pH = 6 -8. Maximal current production obtained at pH = 5 (black), 6 (red), 7 (blue) and 8 (magenta). The error bars represent the standard deviation over 3 independent measurements.

**Fig. S5.**
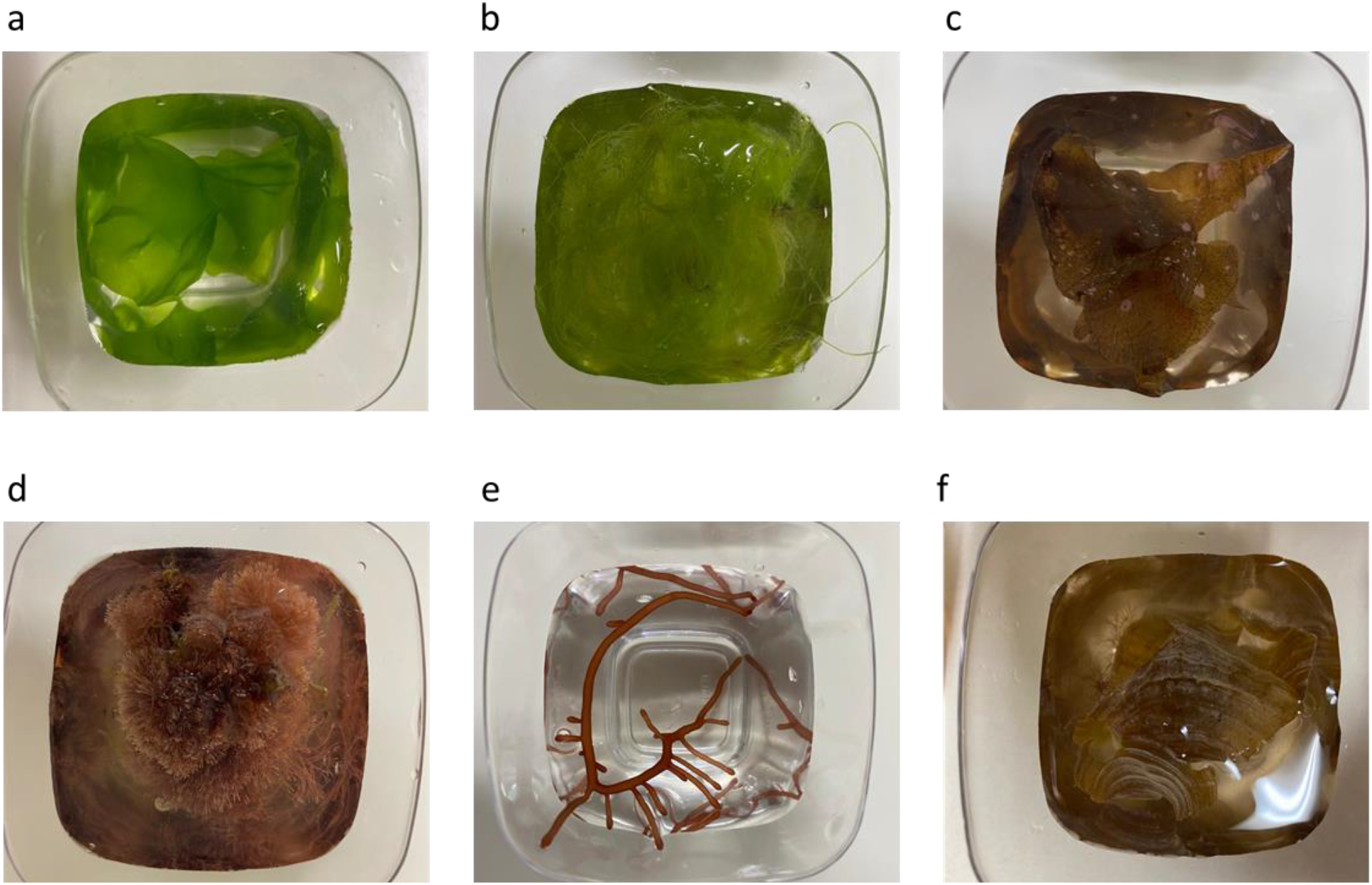
Pictures of the seaweeds from different taxonomic groups. An enlargement of the photos in Fig. S6. **a** *Ulva*. **b** *Cladophora*. **c** *Stypopodium*. **d** *Jania*. **e** *Gracilaria*. **f** *Padina*.

**Fig. S6.**
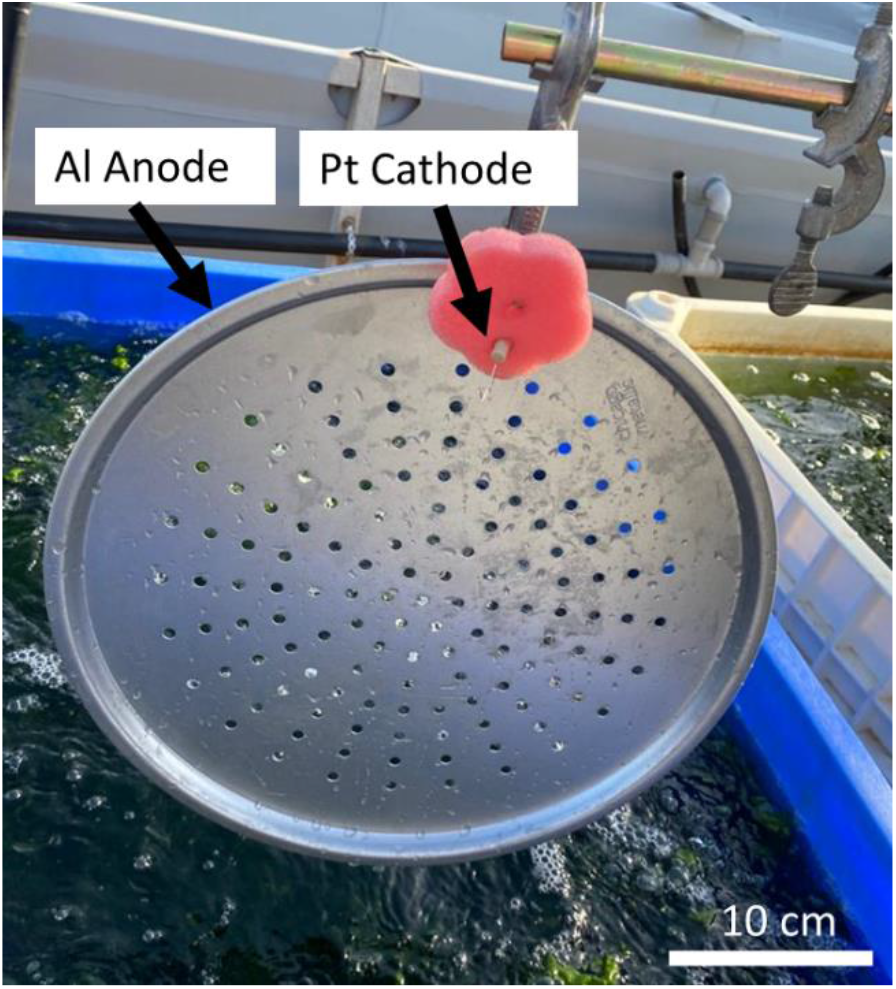
Picture of the electrodes that were used for current production directly from the *Ulva* cultivation pools. In order to make CA measurements directly from the *Ulva* cultivation pool, an Aluminium (Al) round disc was used as Anode and a Platinum wire as cathode. The anode and cathode are marked with black arrows.

**Fig. S7.**
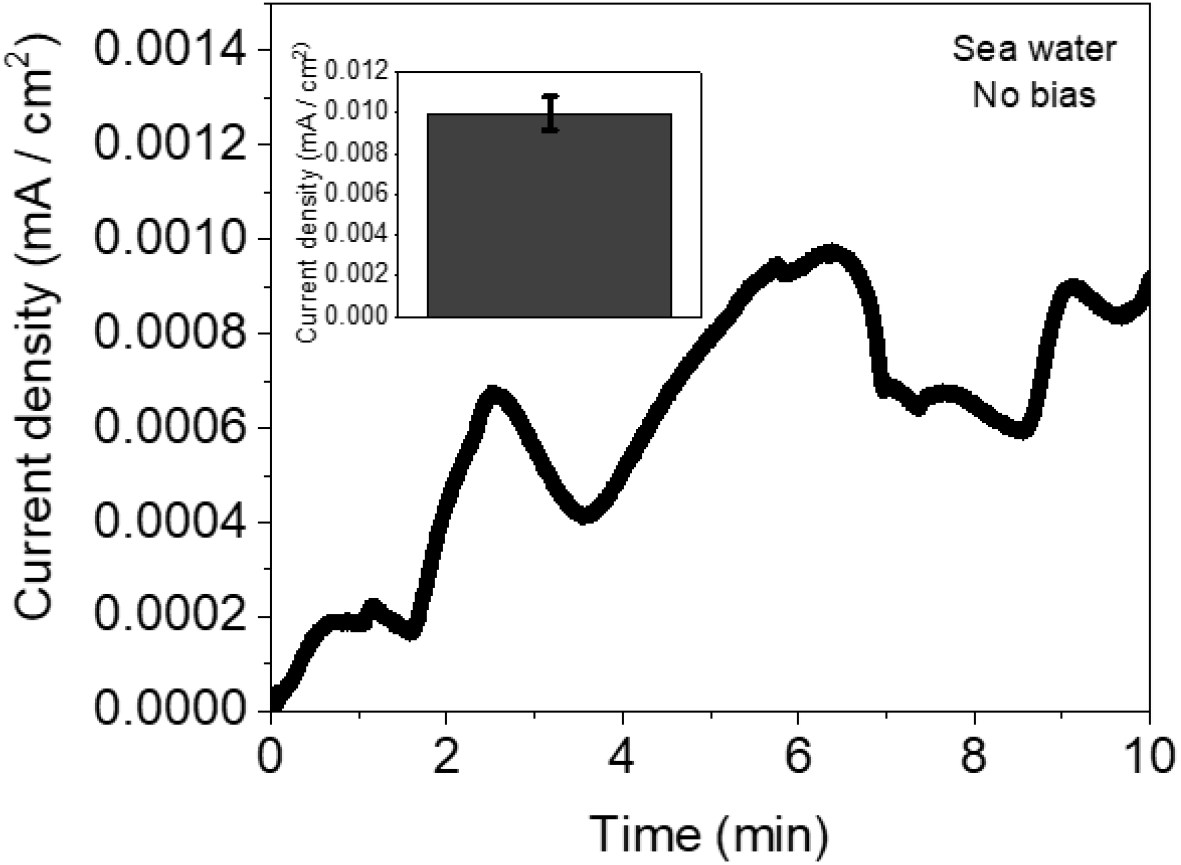
Bias free photocurrent production of *Ulva*. CA of *Ulva* was measured in a 2 electrodes mode in sea water with a stainless-steel clip anode and a Pt wire cathode without application of external potential bias. The error bar in the insert represents the standard deviation over 3 independent measurements.

## References

1. P. Baweja, S. Kumar, D. Sahoo, I. Levine, Chapter 3 - Biology of Seaweeds (Academic Press, San Diego, 2016).

2. H. Dieter, C. Wiencke, K. Bischof, Photosynthesis in Marine Macroalgae (2004), vol. 14.

3. O. G. Mouritsen, Seaweeds: Edible, Available & Sustainable (University of Chicago Press, Chicago & London, ed. 1, 2013).

4. J. A. Raven, C. L. Hurd, Ecophysiology of photosynthesis in macroalgae. Photosynth. Res. 113, 105–125 (2012).

5. M. Aresta, A. Dibenedetto, G. Barberio, Utilization of macro-algae for enhanced CO2 fixation and biofuels production: Development of a computing software for an LCA study. Fuel Process. Technol. 86, 1679–1693 (2005).

6. N. Krupnik et al., Native, invasive and cryptogenic Ulva species from the Israeli Mediterranean sea: risk and potential. Mediterr. Mar. Sci. 19, 132–146 (2018).

7. A. Chemodanov et al., Feasibility study of Ulva sp. (Chlorophyta) intensive cultivation in a coastal area of the eastern Mediterranean sea. Biofuels, Bioprod. Biorefining. 13, 864–877 (2019).

8. A. Qarri, A. Israel, Seasonal biomass production, fermentable saccharification and potential ethanol yields in the marine macroalga Ulva sp. (Chlorophyta). Renew. Energy. 145, 2101–2107 (2020).

9. L. D et al., Role of C 4 carbon fixation in Ulva prolifera, the macroalga responsible for the world’s largest green tides. Commun. Biol. 3, 494 (2020).

10. G. Rosenberg, J. Ramus, Ecological Growth Strategies in the Seaweeds. Mar. Ecol. 8, 233–241 (1982).

11. S. Beer, O. Poryan, C. Larsson, L. Axelsson, Photosynthetic rates of Ulva (chlorophyta) measured by pulse amplitude modulated (pam) fluorometry. Eur. J. Phycol. 35, 69–74 (2000).

12. J. Xu et al., Evidence of coexistence of C3 and C4 photosynthetic pathways in a green-tide-forming alga, Ulva prolifera. PLoS One. 7, 1–10 (2012).

13. T. Beer, A. Israel, Y. Helman, A. Kaplan, Acidification and CO2 production in the boundary layer during photosynthesis in Ulva rigida (Chlorophyta) C. Agardh. Isr. J. Plant Sci. 56, 55–60 (2008).

14. S. Beer, M. Bjork, J. Beardall, Photosynthesis in the Marine Environment (Wiley-Blackwell, 2014).

15. A. Israel, in Photosynthesis in the Marine Environment, S. Beer, M. Bjork, J. Beardall, Eds. (Wiley-Blackwell, 2014; https://www.amazon.com/Photosynthesis-Marine-Environment-Sven-Beer/dp/1119979579), pp. 104–117.

16. M. Ghadiryanfar, K. A. Rosentrater, A. Keyhani, M. Omid, A review of macroalgae production, with potential applications in biofuels and bioenergy. Renew. Sustain. Energy Rev. 54, 473–481 (2016).

17. N. Gaurav, S. Sivasankari, G. S. Kiran, A. Ninawe, J. Selvin, Utilization of bioresources for sustainable biofuels: A Review. Renew. Sustain. Energy Rev. 73, 205– 214 (2017).

18. M. Zollmann, H. Traugott, A. Chemodanov, A. Liberzon, A. Golberg, Exergy efficiency of solar energy conversion to biomass of green macroalgae Ulva (Chlorophyta) in the photobioreactor. Energy Convers. Manag. 167, 125–133 (2018).

19. X. Wei, H. Lee, S. Choi, Biopower generation in a microfluidic bio-solar panel. 228, 151–155 (2016).

20. A. Rhoads, H. Beyenal, Z. Lewandowski, Microbial Fuel Cell using Anaerobic Respiration as an Anodic Reaction and Biomineralized Manganese as a Cathodic Reactant. Environ. Sci. Technol. 39, 4666–4671 (2005).

21. J. Menicucci et al., Procedure for Determining Maximum Sustainable Power Generated by Microbial Fuel Cells. Environ. Sci. Technol. 40, 1062–1068 (2006).

22. K. Rabaey, N. Boon, M. Höfte, W. Verstraete, Microbial Phenazine Production Enhances Electron Transfer in Biofuel Cells. Environ. Sci. Technol. 39, 3401–3408 (2005).

23. B. R. Ringeisen et al., High Power Density from a Miniature Microbial Fuel Cell Using Shewanella oneidensis DSP10. Environ. Sci. Technol. 40, 2629–2634 (2006).

24. B. Min, S. Cheng, B. E. Logan, Electricity generation using membrane and salt bridge microbial fuel cells. Water Res. 39, 1675–1686 (2005).

25. D. R. Bond, D. R. Lovley, Electricity Production by Geobacter sulfurreducens Attached to Electrodes. Appl. Environ. Microbiol. 69, 1548–1555 (2003).

26. X. Fang, S. Kalathil, E. Reisner, Semi-biological approaches to solar-to-chemical conversion (Royal Society of Chemistry, 2020; https://pubmed.ncbi.nlm.nih.gov/32538416/), vol. 49.

27. A. Bergel, D. Féron, A. Mollica, Catalysis of oxygen reduction in PEM fuel cell by seawater biofilm. Electrochem. commun. 7, 900–904 (2005).

28. B. K. Kaiser et al., Fatty aldehydes in cyanobacteria are a metabolically flexible precursor for a diversity of biofuel products. PLoS One. 8, e58307 (2013).

29. A. J. McCormick et al., Biophotovoltaics: oxygenic photosynthetic organisms in the world of bioelectrochemical systems. Energy Environ. Sci. 8, 1092–1109 (2015).

30. R. I. Pinhassi et al., Hybrid bio-photo-electro-chemical cells for solar water splitting. Nat. Commun. 7, 1–10 (2016).

31. F. Zhao et al., Light Induced H Evolution from a Biophotocathode Based on Photosystem 1 - Pt Nanoparticles Complexes Integrated in Solvated Redox Polymers Films. J Phys Chem B. 119, 13726–13731 (2015).

32. A. Efrati et al., Assembly of photo-bioelectrochemical cells using photosystem I-functionalized electrodes. Nat. Energy. 1, 15021 (2016).

33. E. A. Gizzie et al., Photosystem I-polyaniline/TiO 2 solid-state solar cells: simple devices for biohybrid solar energy conversion. 8, 3572–3576 (2015).

34. N. Sekar, R. Jain, Y. Yan, R. P. Ramasamy, Enhanced photo-bioelectrochemical energy conversion by genetically engineered cyanobacteria. Biotechnol. Bioeng. 113, 675–679 (2016).

35. M. Sawa, Electricity generation from digitally printed cyanobacteria. Nat. Commun. 8 (2017), doi:10.1038/s41467-017-01084-4.

36. K. P. Sokol et al., Bias-free photoelectrochemical water splitting with photosystem II on a dye-sensitized photoanode wired to hydrogenase. Nat. Energy. 3, 944–951 (2018).

37. G. Saper et al., Live cyanobacteria produce photocurrent and hydrogen using both the respiratory and photosynthetic systems. Nat. Commun. 9, 2168 (2018).

38. P. Bombelli et al., Quantitative analysis of the factors limiting solar power transduction by Synechocystis sp. PCC 6803 in biological photovoltaic devices. Energy Environ. Sci. 4, 4690–4698 (2011).

39. A. Cereda et al., A bioelectrochemical approach to characterize extracellular electron transfer by Synechocystis sp. PCC6803. PLoS One. 9, 91484 (2014).

40. Y. Shlosberg et al., NADPH performs mediated electron transfer in cyanobacterial-driven bio-photoelectrochemical cells. iScience. 24, 101892 (2021).

41. Y. Zou, J. Pisciotta, R. B. Billmyre, I. V Baskakov, Photosynthetic microbial fuel cells with positive light response. Biotechnol. Bioeng. 104, 939–946 (2009).

42. A. C. Gonzalez-Aravena, K. Yunus, L. Zhang, B. Norling, A. C. Fisher, Tapping into cyanobacteria electron transfer for higher exoelectrogenic activity by imposing iron limited growth. RSC Adv. 8, 20263–20274 (2018).

43. B. E. Logan, Exoelectrogenic bacteria that power microbial fuel cells. Nat. Rev. Microbiol. 7, 375–381 (2009).

44. R. I. Pinhassi et al., Photosynthetic membranes of Synechocystis or plants convert sunlight to photocurrent through different pathways due to different architectures. PLoS One. 10, e0122616 (2015).

45. W. Zhang, X. Chen, Y. Wang, L. Wu, Y. Hu, Experimental and modeling of conductivity for electrolyte solution systems. ACS Omega. 5, 22465–22474 (2020).

46. J. Tschörtner, B. Lai, J. O. Krömer, Biophotovoltaics: Green power generation from sunlight and water. Front. Microbiol. 10, 866 (2019).

47. J. Huang, B. Sun, X. Zhang, Electricity generation at high ionic strength in microbial fuel cell by a newly isolated Shewanella marisflavi EP1. Appl. Microbiol. Biotechnol. 2009 854. 85, 1141–1149 (2009).

48. † Hong Liu, † and Shaoan Cheng, †,‡ Bruce E. Logan*, Power Generation in Fed-Batch Microbial Fuel Cells as a Function of Ionic Strength, Temperature, and Reactor Configuration. Environ. Sci. Technol. 39, 5488–5493 (2005).

49. L. R. Dartnell, M. C. Storrie-Lombardi, J. M. Ward, Complete fluorescent fingerprints of extremophilic and photosynthetic microbes. Int. J. Astrobiol. 9, 245–257 (2010).

50. Y. Bi, Z. Zhou, Absorption and transport of inorganic carbon in kelps with emphasis on Saccharina japonica. Appl. Photosynth. - New Prog. (2016).

51. A. J. McCormick et al., Photosynthetic biofilms in pure culture harness solar energy in a mediatorless bio-photovoltaic cell (BPV) system. Energy Environ. Sci. 4, 4699–4709 (2011).

52. P. Bombelli, T. Müller, T. W. Herling, C. J. Howe, T. P. J. Knowles, A high power-density, mediator-free, microfluidic biophotovoltaic device for cyanobacterial cells. Adv. Energy Mater. 5 (2015), doi:10.1002/aenm.201401299.

53. H. Ochiai, H. Shibata, Y. Sawa, M. Shoga, S. Ohta, Properties of semiconductor electrodes coated with living films of cyanobacteria. Appl. Biochem. Biotechnol. 8, 289–303 (1983).

54. J. C.-W. Lan, K. Raman, C.-M. Huang, C.-M. Chang, The impact of monochromatic blue and red LED light upon performance of photo microbial fuel cells (PMFCs) using Chlamydomonas reinhardtii transformation F5 as biocatalyst. Biochem. Eng. J. 78, 39–43 (2013).

55. A. Laohavisit et al., Enhancing plasma membrane NADPH oxidase activity increases current output by diatoms in biophotovoltaic devices. Algal Res. 12, 91–98 (2015).

56. N. Krupnik et al., Dust-borne microbes affect Ulva ohnoi’s growth and physiological state. FEMS Microbiol. Ecol. 97, fiab020 (2021).

57. Y. Lehahn, K. N. Ingle, A. Golberg, Global potential of offshore and shallow waters macroalgal biorefineries to provide for food, chemicals and energy: Feasibility and sustainability. Algal Res. 17, 150–160 (2016).

58. F. J. Rawson, A. J. Downard, K. H. Baronian, Electrochemical detection of intracellular and cell membrane redox systems in Saccharomyces cerevisiae. Sci. Rep. 4, 1–9 (2014).

59. S. Shiva Kumar, V. Himabindu, Hydrogen production by PEM water electrolysis – A review. Mater. Sci. Energy Technol. 2, 442–454 (2019).

60. A. J. Lawaetz, C. A. Stedmon, Fluorescence intensity calibration using the Raman scatter peak of water. Appl. Spectrosc. 63, 936–940 (2009).

